# SLC26A2-mediated sulfate metabolism is essential for the tooth development

**DOI:** 10.1101/2023.03.30.534919

**Authors:** Y. Yoshida, T. Inubushi, M. Yokoyama, P. Nag, A. Oka, T. Murotani, R. Kani, Y. Shiraishi, H. Kurosaka, Y. Takahata, R. Nishimura, P. Papagerakis, T. Yamashiro

## Abstract

The sulfate transporter gene SLC26A2 is responsible for diastrophic dysplasia, which represents skeletal dysplasia in humans. This highlights the importance of sulfate metabolism in skeletal formation. SLC26A2-related chondrodysplasia is also known to exhibit abnormalities in craniofacial and tooth development. Although the function of SLC26A2 in mammals has been investigated using genetic mouse models, the essential role of SLC26A2 during craniofacial and tooth development has not been elucidated.

In this study, we demonstrate the pivotal roles of SLC26A2-mediated sulfate metabolism during tooth development. Analysis of *Slc26a2* expression reveals that it is predominantly expressed in dental tissues, including odontoblasts and ameloblasts, during tooth development. *Slc26a2* knockout mice (*Slc26a2-KO-Δexon2*) exhibit a retrognathic upper jaw, small upper incisors, and hypoplasia of upper molars. Additionally, upper incisors and molars in *Slc26a2-KO-Δexon2* mice display flattened odontoblasts and nuclei that lack intracellular polarity. In contrast, tooth phenotype is not remarkable in lower incisors and molars.

Furthermore, the expression of odontoblast differentiation markers, *Dspp* and *Dmp1*, is significantly decreased in the upper molars of *Slc26a2*-deficient mice. *Ex vivo* organ culture of tooth germs by implantation of *Slc26a2*-deficient tooth germs under the kidney capsule reveals hypoplasia of the dentin matrix as well as tooth root shortening. *In vitro* studies using human dental pulp stem cells (hDPSCs) show that the expression levels of *Dspp* and *Dmp1* in *shSlc26a2* knockdown cells are significantly decreased compared to control cells.

Collectively, our data demonstrate that SLC26A2-mediated sulfate metabolism is essential for tooth development. This study may provide insight into the mechanisms underlying tooth abnormalities in patients with recessively inherited chondrodysplasias caused by mutations in the SLC26A2 gene.

## Introduction

A number of different congenital craniofacial anomalies and tooth developmental defects have been reported, including cleft lip and/or palate. Most congenital defects are caused by genetic or environmental factors or the combination thereof. Genetic research using genetic mice models has revealed several transcription factors and signaling molecules, including WNTs, BMPs, and FGFs have been identified as genes in human isolated and syndromic odontogenesis. The environmental causes of congenital anomalies include fetal infection, maternal illness, nutritional deficiencies, drug ingestion, chemical exposures, air pollution, and radiation (Baldacci et al., 2018). However, the etiology of congenital anomalies has not yet been fully elucidated.

Sulfate or sulfate ion (SO_4_^2-^) is an important anion involved in a wide range of significant biological processes, including biosynthesis and metabolism of a variety of endogenous biological molecules, and is essential for cell growth and development of the organism (Foster and Mueller, 2018; Soares da Costa et al., 2017). Since sulfate cannot freely pass through the plasma membranes of cells, transport mechanisms are required for the movement of sulfate into and out of mammalian cells (Alper and Sharma, 2013). Such mechanisms are also necessary for sulfate absorption from the gastrointestinal tract and re-absorption by the renal tubules (Seidler and Nikolovska, 2019). After sulfate is transported into the cytoplasm or nucleus, 3’-phosphoadenosine 5’-phosphosulfate (PAPS) is synthesized from ATP and sulfate by two enzymatic reactions: ATP sulfurylase and adenosine 5’-phosphosulfate kinase (APS kinase). After PAPS is transported to the Golgi apparatus, sulfotransferases transfer sulfate groups from PAPS to glycosaminoglycans and tyrosine. In addition, cytosolic sulfotransferases transfer sulfate groups from PAPS to steroid hormones in cytosol (Klaassen and Boles, 1997). However, part of the sulfate supply is also known to be derived from the breakdown of the sulfur-containing amino acids cysteine and methionine (Elgavish and Meezan, 1991; Markovich and Aronson, 2007).

In mammals, the SLC26 gene family, which encodes anion transporters, consists of 11 genes: SLC26A1-A11. The SLC26 gene family transports a broad range of anionic substrates, including sulfate, HCO^3−^, Cl^−^, oxalate, I^−^, and formate. SLC26A1 and SLC26A2 encode the proteins SAT1 and DTDST, respectively, which are sulfate/chloride exchangers that function as cell membrane sulfate transporters and enable the intracellular transport of inorganic sulfate (Kere, 2006). In humans, mutations in the SLC26A2 gene cause a spectrum of recessively inherited chondrodysplasias. Although the phenotype differs according to the type of *SLC26A2* mutation, the main clinical features are a short stature, joint contractures, club feet, shortening of limbs, and a waddling gait (Cai et al., 2015). Recent reports also suggest that *SLC26A2* mutations are associated with a wide range of clinical manifestations in the craniomaxillofacial region, including a large upper facial height, micrognathia, high palate, cleft palate (25%-60%), congenital anodontia (30%), and dwarf teeth (Härkönen et al., 2021). Mutations in the other eight members of the SLC26 family have also been implicated in human disease. However, unlike SLC26a2 mutations, mutations in the other SLC26 family members do not induce any abnormalities in the skeletal system or craniofacial region.

The function of *Slc26a2* in mammals has been investigated using genetic mice models. Forlino et al. generated a Dtdst knock-in mouse model harboring human mutations, and the mice showed a partial loss of function of the sulfate transporter. These mice exhibited a short stature, joint contracture, reduced toluidine blue staining of cartilage, and irregular chondrocyte size (Forlino et al., 2005). Zheng et al. generated Slc26a2^-/-^ mice to investigate the effects of SLC26A2 deficiency in chondrodysplasia (Zheng et al., 2019). Although patients with mutations in the SLC26A2 gene reportedly show abnormalities in the craniomaxillofacial region, including dwarf teeth and congenital absence of teeth, the essential role of SLC26A2 during tooth development has not been elucidated.

In the present study, we investigate the expression pattern of Slc26a2 in developing tooth germs, perform morphological and histological evaluations of tooth germs in Slc26a2 knockout mice, and examine the effects of Slc26a2 deficiency on enamel and dentin matrix formation using kidney capsule grafting. This is the first paper to demonstrate the pivotal role of the SLC26A2-mediated sulfate transporter during tooth development.

## Materials and methods

### Animal

Pronuclear stage embryos from C57BL6/J mice were purchased from ARK Resource (Kumamoto, Japan). Recombinant Cas9 protein, crRNA, and tracrRNA were obtained from Integrated DNA technology. For a generation of Slc26a2-KO-△exon2 mice, we used the Technique for Animal Knockout System by Electroporation (TAKE) as previously described (Inubushi T., 2022). For more information, please see **the appendix**.

### Tissue preparation, histology and *in situ* hybridization

Maxillary and mandibular from control and *Slc26a2-KO-△exon2* mice at P0 were collected and fixed in 4% paraformaldehyde. A mild decalcifier, osteosoft (sigma-aldrich), was used for tooth decalcification. Sagittal sections of paraffin embedded mandible were prepared and used for haematoxylin & eosin (H&E) and *in situ* hybridization as previously described (Sarper et al., 2018). For more information, please see **the appendix**.

### Immunohistochemistry

Frozen section of 12 µm thickness were used for immunostaining and incubated with M.O.M (mouse on mouse) blocking reagent, 5% goat serum/PBS, and 0.1% sodium citrate buffer. The primary and secondary antibodies used in this study were described in **the appendix**.

### Laser microdissection

We performed Laser microdissection as previously reported(Sarper et al., 2018). For more information, please see **the appendix**.

### RNA extraction and qRT-PCR analysis

The protocol for RNA extraction and qPCR analysis was as reported previously(Inubushi et al. 2012). For more information, please see **the appendix**.

### Transplantation Under the Kidney Capsule of Mice

Control and Slc26a2-KO E18.5 upper molar tooth germ were implanted under the renal capsule of 8-week-old BALB/cSlc-nu/nu mice. 4 weeks later, the implanted tooth embryos were harvested and micro-CT was performed. 50-µm sagittal slice images were taken with VG Studio and the image processing software The roots were cut in ImageJ and the volume of the crown dentin was measured.

### Blyscan sulfated glycosaminoglycan assay

The Blyscan sulfated glycosaminoglycan assay kit (Biocolor, Carrickfergus, United Kingdom) quantitatively measures sulfated proteoglycans and GAGs in biological samples. The assay was performed according to the manufacturer’s instructions. For more information, please see **the appendix.**

**For Ethics statement, Whole-mount skeletal staining, Micro-Computed Tomography (MicroCT), Reanalysis of public scRNA-seq data, Cell culture and lentivirus transduction, Alcian blue staining, Statistical analysis** See **the appendix**.

## Results

### Slc26a2 is predominantly expressed among sulfate transporter family genes during tooth development

To understand the role of SLC26a2 in tooth development, we quantitatively analyzed the expression of sulfate transporter family genes, including *Slc26a1* and *Slc26a2*, in developing tooth germ. We reanalyzed the public data set of single-cell RNA sequencing on the isolated mice incisors at P0. We identified 11 cell population clusters of odontoblasts, sub-odontoblasts, dental mesenchyme 1 and 2, ameloblasts, pre-ameloblasts, inner enamel epithelium or outer enamel epithelium (IEE/OEE), stratum intermedium or stellate reticulum (SI/SR), leukocytes, erythrocytes, and endothelial cells (Fig. 1A and Appendix Figure 1). We next characterized the expression of *Slc26a1, Slc26a2*, *Slc26a6*, *Slc26a7*, *Slc26a10*, and *Slc26a11* in the data sets. A dot plot showed that *Slc26a2* was more strongly expressed in all clusters, including ameloblast and odontoblast clusters, than other sulfate transporter family genes (Fig. 1B).

We next used *in situ* hybridization to confirm the expression pattern of *Slc26a2* in tooth development in mice at embryonic day (E) 18.5. The prominent expression of *Slc26a2* was observed in odontoblasts and ameloblasts at E18.5 (Fig. 1C). Interestingly, at each developmental stage, the expression of *Slc26a2* was found to be markedly higher than that of *Slc26a1* (Appendix Figure 2A). Furthermore, the expression of *Slc26a2* increased with developmental stage, with craniofacial formation beginning at E9.5 and the tooth bud forming at E13.5 (Appendix Figure 2A). At E18.5, the expression of *Slc26a2* was higher than that of *Slc26a1* in the upper molars (Appendix Figure 2B). These results suggest that *Slc26a2* plays a major role in the transport of sulfate ions through the cell membrane during tooth development.

**Figure 1.**
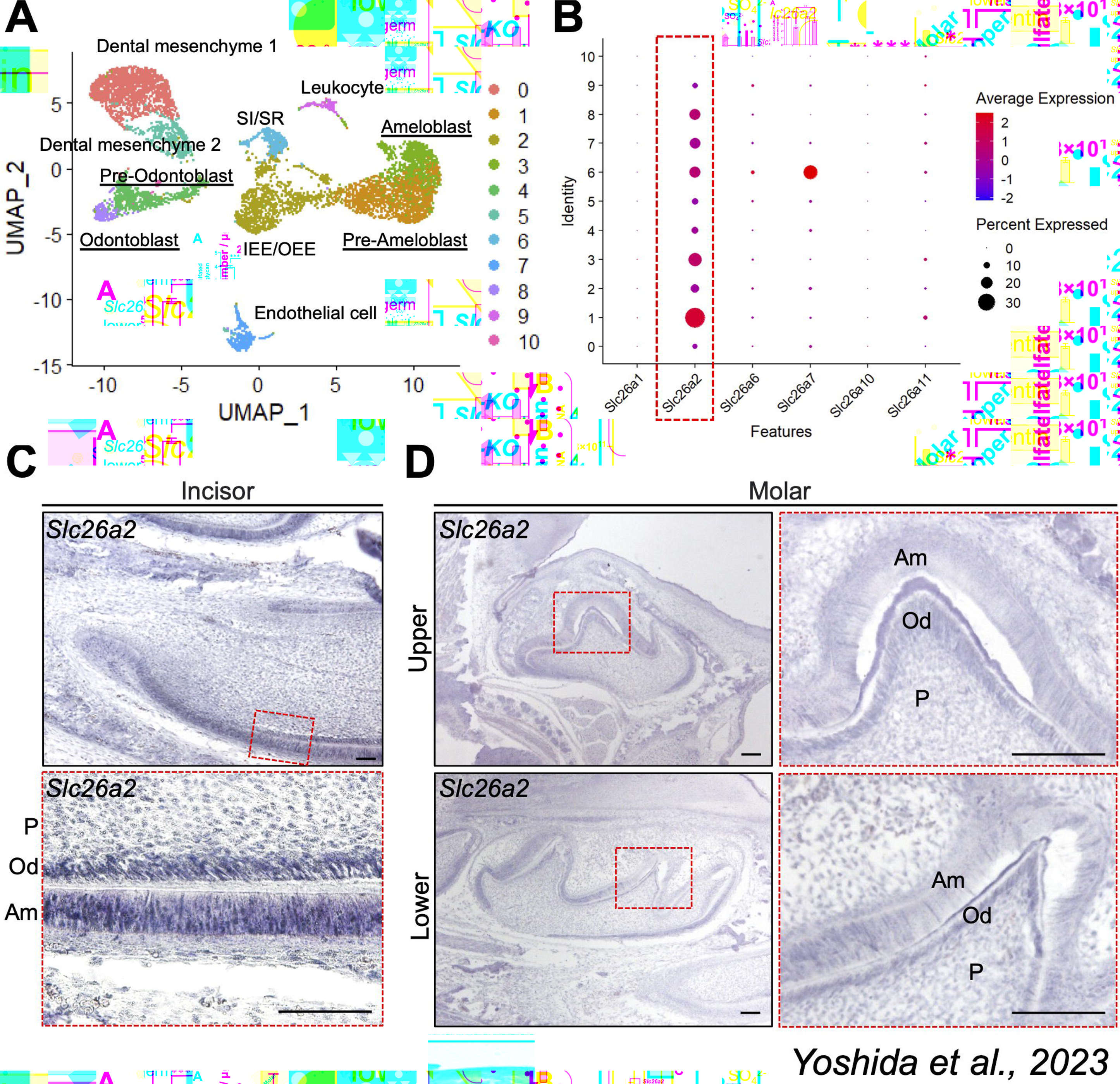
Expression pattern of *Slc26a2* in tooth development in mice. (A) Re-analysis of the public data set of single-cell RNA sequencing on the isolated mice incisors at P0. Ten cell population clusters, including odontoblasts (cluster 8), pre-odontoblasts (cluster 4), ameloblasts (cluster 3), and pre-ameloblasts (cluster 1), are identified in mice tooth germs at P0. (B) Dot plot showing the expression of SLC26 family genes in various clusters. Dot size represents the % of cells expressing a specific gene. The intensity of color indicates the average expression level for the gene in the cluster. *Slc26a2* is predominantly expressed in mice tooth germs. (C) The expression pattern of *Slc26a2* during tooth development at E18.5. Frontal sections of the upper molar, sagittal sections of the lower incisor, and molar in wild type mice were processed by RNA *in situ hybridization*. The boxed area by the red dotted line is magnified and shown in adjacent image. am: ameloblast; od: odontoblasts; P: pulp. Scale bars: 100 μm. All the samples are wild-type C57B6/J.

**Figure 2.**
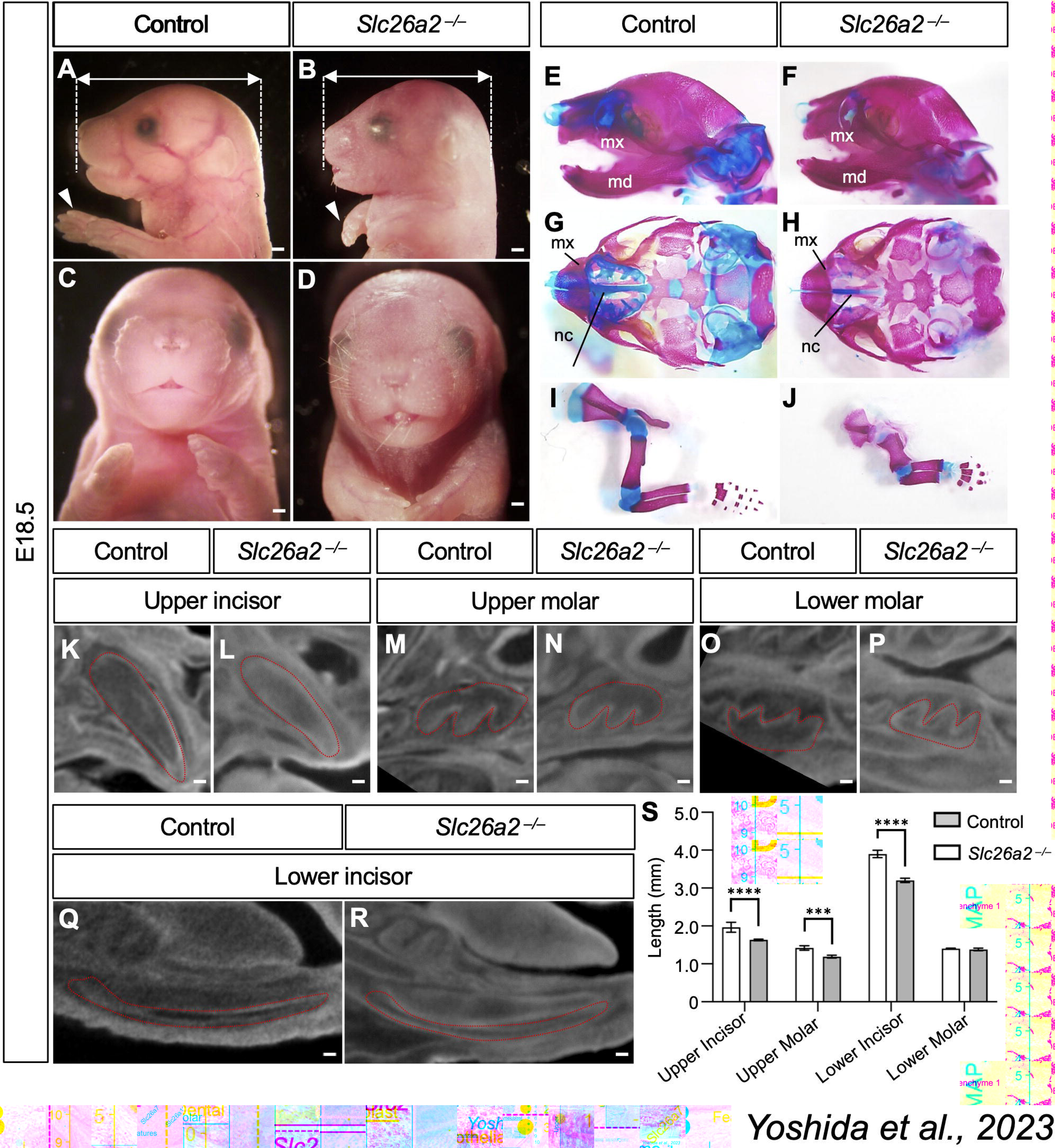
*Slc26a2-KO-△exon2* mice show skeletal and dental abnormality. (A-D) Gross phenotype of *Slc26a2-KO-△exon2* mice at E 18.5. Whole mount images of *Slc26a2-KO-△exon2* revealed hypoplasia of the maxilla and very short limbs (white arrows). Lateral (A, B) and frontal (C, D) views of the head. (E-J) Whole-mount skeletal preparations of craniofacial bones (E, F), cranial base (G, H), and limbs (I, J) in control and *Slc26a2-KO-△exon2* mice at E 18.5. Whole-mount skeletal preparations show that *Slc26a2-KO-△exon2 mice* had chondrodysplasia and reduced Alcian blue staining of the limbs and cranial cartilage. mx: maxilla; md: mandible; nc: nasal cavity. Scale bars: 2 mm. (K-R) Contrast-enhanced Micro-CT images of control and *Slc26a2-KO-△exon2* mice tooth germs at E18.5. *Slc26a2-KO-△exon2* mice show smaller width in the upper molars and incisors than the control mice. Scale bar: 50 μm. (S) Quantitative assessment of the upper and lower tooth size in control and *Slc26a2-KO-△exon2* mice at E 18.5. *Slc26a2-KO-△exon2* mice show hypoplasia of the upper incisor and molars, and lower incisors, but not the lower molars comparing to the control mice. Means ± s.d. (n = 3) are shown as horizontal bars. *P* values were determined by one-way ANOVA. \**P* < 0.01.

### Slc26a2-deficient mice show retrognathic upper jaw and hypoplasia of the upper teeth

To investigate the role of *Slc26a2* deficiency in tooth development, we generated *Slc26a2* knockout mice by targeted deletion of *Slc26a2* exon 2 (*Slc26a2-KO-△exon2*) using the CRISPR-Cas9 gene editing method (Makino et al., 2016). Previous reports showed that *Slc26a2*-*KO* mice died immediately after birth, with no respiratory movement and an overall skeletal phenotype characterized by a short neck, small chest, and very short limbs (Zheng et al., 2019). Consistently, *Slc26a2-KO-△exon2* mice died immediately after birth due to respiratory abnormalities. Whole mount images of *Slc26a2-KO-△exon2* revealed hypoplasia of the maxilla rather than the mandibula (Fig. 2A-D). Micro-computed tomography (CT) of *Slc26a2-KO-△exon2* mice demonstrated a short stature, small chest, and very short limbs. The long tubular bone was shorter in length and longer in diameter than the control mice (Appendix Figure 3A, B). Whole-mount skeletal preparations showed that *Slc26a2-KO-△exon2* had chondrodysplasia and reduced Alcian blue staining of the cartilage (Appendix Figure 3C, D). In the cranio-maxillofacial region, *Slc26a2-KO-△exon2* mice showed retrognathic upper jaw, hypoplasia of nasal cartilage (Fig. 2E, F), small cranial base (Fig. 2G, H), and short ribs (Fig. 2I–J). At E18.5, mutant and control tooth germs were examined by contrast-enhanced micro-CT (Fig. 2K–R). *Slc26a2-KO-△exon2* mice showed a shorter long diameter from the cervical loop to the incisal margin of the upper incisor tooth germ than the control mice. Similarly, the upper molar tooth germs in *Slc26a2-KO-Δexon2* mice were shorter with regard to both crown proximal-central width and height compared to control mice. In contrast, there were no significant differences in the width or height of the lower incisors and lower molars (Fig. 2S).

**Figure 3.**
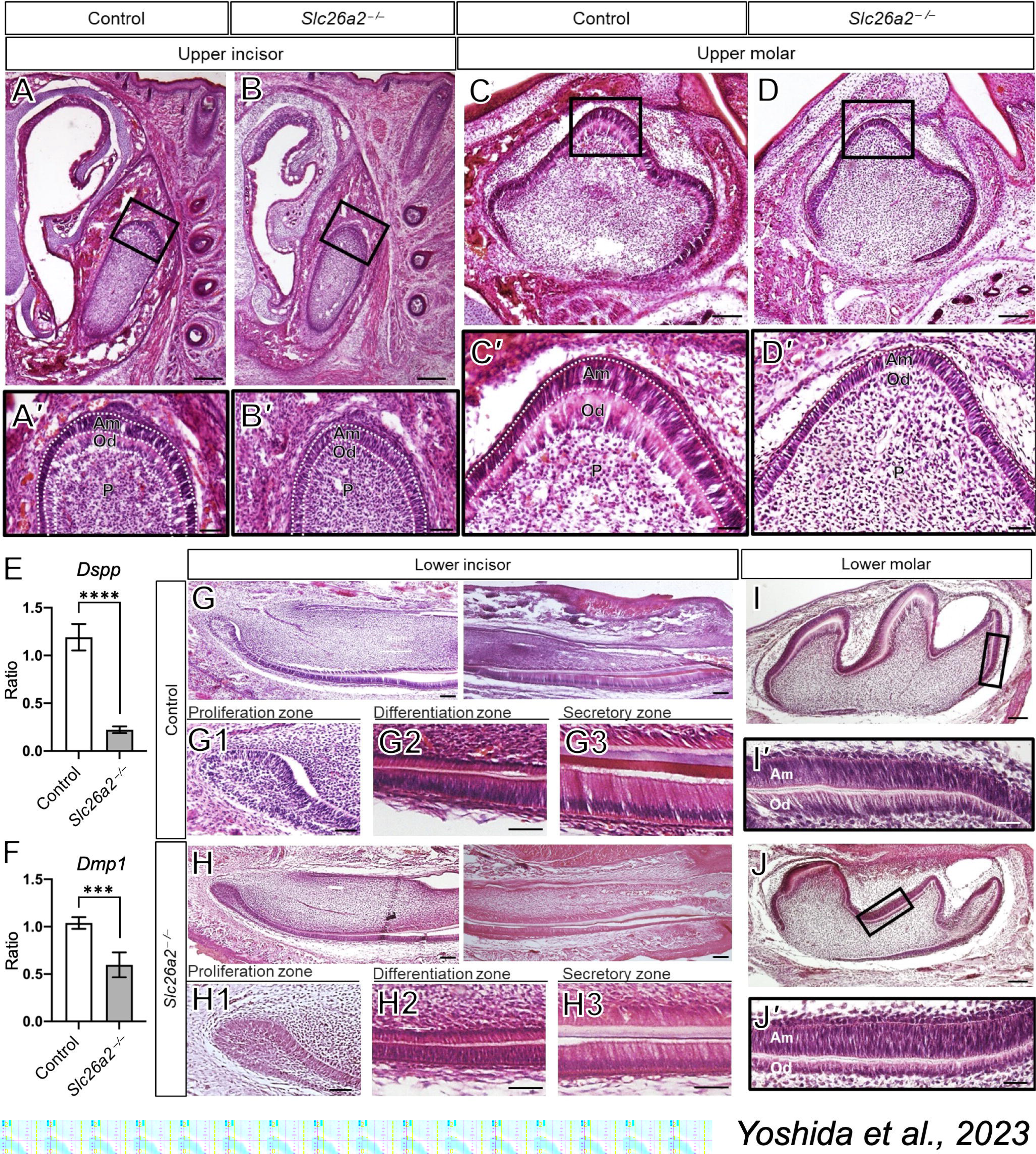
*Slc26a2* deficiency leads to impaired differentiation of odontoblasts in the upper tooth. (A-D) H&E staining of frontal sections of upper incisor and molar tooth germ. Upper incisor and molar tooth germ in *Slc26a2-KO-△exon2* mice show flattened odontoblasts cell body and nuclei, and loss of intracellular polarity. (A’-D’) Areas indicated by the black line are enlarged in lower panels. (E, F) qPCR analysis of *Dspp* and *Dmp1* expression in odontoblast of upper tooth germ at E18.5. In *Slc26a2-KO-△exon2* mice tooth germ, the expression of *Dspp* and *Dmp1* in upper molar tooth germ was significantly decreased compared to the control tooth germ. Means ± s.d. (n=3) are shown as horizontal bars. *P* values were determined by unpaired Student’s *t*-test. \**P* < 0.01. Dentin sialophosphoprotein: *Dspp*; Dentin matrix protein 1: *Dmp1*. (G-H) H&E staining of sagittal sections of lower incisor and molar tooth germ. The overall structure of lower incisor and molar tooth germ is comparable in control and *Slc26a2-KO-△exon2* mice. High magnification images corresponding to the area of the proliferation zone (G1 in control mice and H1 in *Slc26a2-KO-△exon2* mice), the differentiation zone (G2 in control mice and H2 in *Slc26a2-KO-△exon2* mice), and the secretary zone (G3 in control mice and H3 in *Slc26a2-KO-△ exon2* mice). In the differentiation zone, the pre-secretary ameloblasts layer is larger in *Slc26a2-KO-△exon2* than in control mice. The height of secretary and mature ameloblasts are lower in *Slc26a2-KO-△exon2* than in control mice. The reduction of enamel and dentin thickness is observed in the incisor tooth germ of *Slc26a2-KO-△exon2* mice. The tooth phenotype in lower molar tooth germ in *Slc26a2-KO-△exon2* mice is much milder than upper molar tooth germ. Am: ameloblast; PD: pre-dentin; D: dentin; E: enamel; Od: odontoblasts; P: dental pulp. Scale bars, 200 μm in A-B; 100 μm in C-D, G-I; 50 μm in A’-D’, G2-G3, H2-H3, I’-J’; 20 μm in G1, G2.

### Slc26a2 deficiency leads to the impaired differentiation of odontoblasts and ameloblasts

We performed a histological evaluation to clarify the effects of *Slc26a2* deficiency on tooth development. Upper incisors and molars in *Slc26a2-KO-△exon2* mice show flattened odontoblasts and nuclei that do not show intracellular polarity (Fig. 3A–D, A’-D’). Ameloblasts of the mutants were not affected.

To further examine the effect of *Slc26a2* deficiency on odontoblast differentiation, we evaluated the expression of the odontoblast marker genes Dentin sialolin protein (*Dspp*) and Dentin matrix protein 1 (*Dmp1*), which code non-collagenous organic substances in dentin and show an increased expression in association with odontoblast differentiation (Chen et al., 2016) (Yamashiro et al., 2007). In *Slc26a2-KO-△exon2* mice tooth germ, the expression of *Dspp* and *Dmp1* of upper molars was significantly decreased compared to the control group (Fig. 3E, F). The tooth phenotype in *Slc26a2-KO-△exon2* mice is much milder in lower incisor and molar tooth germs than in upper incisor and molar tooth germs (Fig. 3G–J,G1-G3,H1-H3,I’.J’). Taken together, these results demonstrate that *Slc26a2* is required for odontoblast differentiation of upper incisors and molars in mice.

### Slc26a2-deficient human dental pulp stem cells show defective differentiation into odontoblasts

To gain further insight into the effect of *Slc26a2* knockdown on the differentiation of dental pulp stem cells into odontoblasts, human dental pulp stem cells (hDPSCs) were used to analyze the odontoblast differentiation potential. *Slc26a2* knockdown cells were generated via lentivirus-mediated delivery of shRNA, and the expression of *Slc26a2* was decreased by more than 80% in *shSlc26a2* knockdown cells (Fig. 4A). In *shSlc26a2* knockdown cells, the expression of *Dspp* and *Dmp1* was significantly decreased compared to the control cells (Fig. 4A). This result indicates that *Slc26a2* knockdown can directly affect the differentiation of pulp stem cells into odontoblasts.

**Figure 4.**
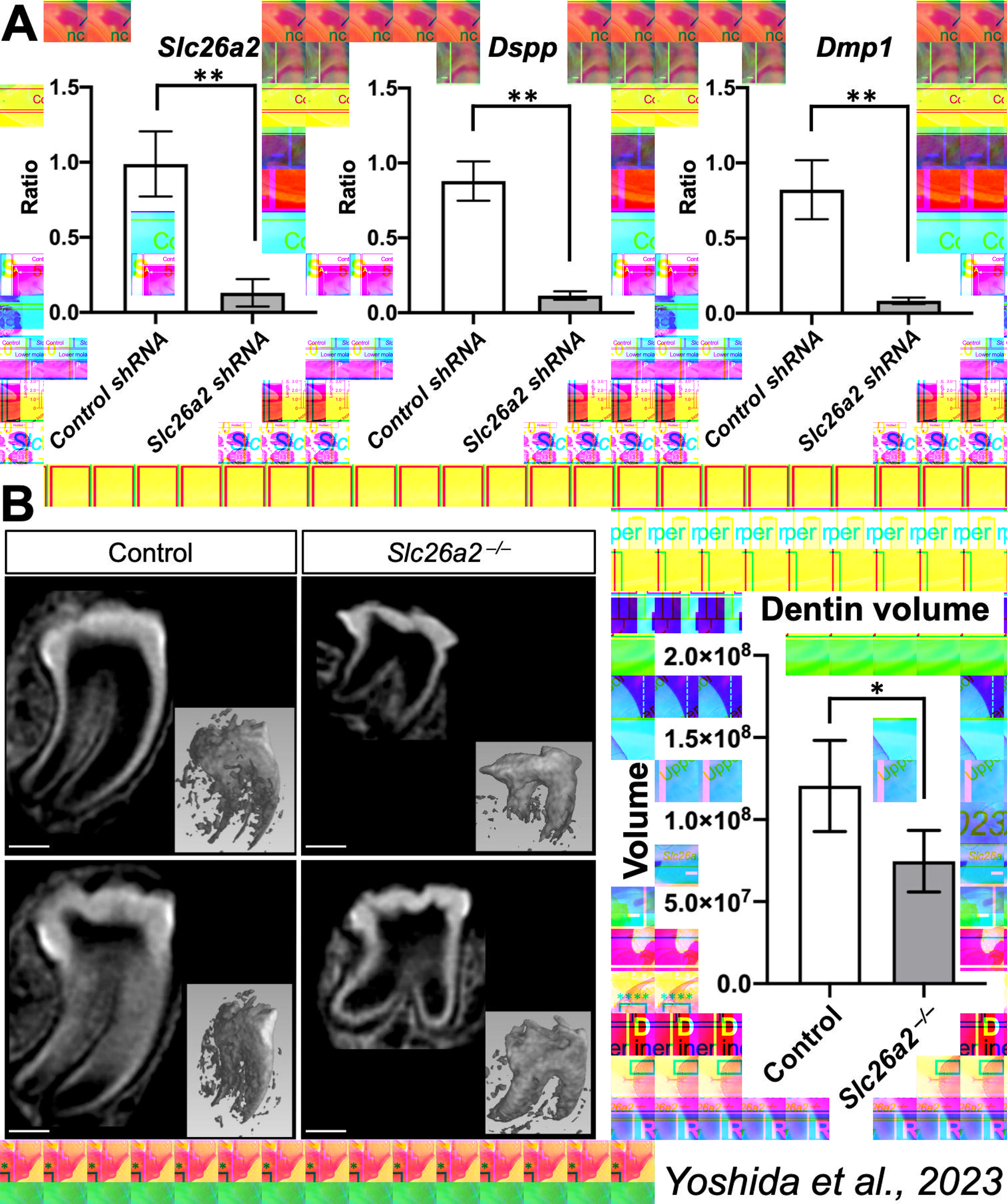
Analysis of the effect of *Slc26a2* deletion in dentinogenesis. (A) *Slc26a2*-depleted and control DPSCs were cultured for 7 days in odontoblasts differentiation medium. The expression of *Slc26a2, Dspp* and *Dmp1* was evaluated by qPCR. Note that *Slc26a2, Dspp* and *Dmp1* expression was significantly decreased in *Slc26a2*-depleted cells compared to the control cells. Gapdh was used as an internal control for normalization. Means ± s.d. (n=3) are shown as horizontal bars. *P* values were determined by unpaired Student’s *t*-test. \*\**P* < 0.01. (B) *Ex vivo* organ culture of tooth germ by implantation of the tooth germ under the kidney capsule of nude mice. After 4 weeks of organ culture, the implanted tooth germs were collected and analyzed by micro-CT. The sagittal sections of control and *Slc26a2-KO-△exon2* mice tooth germ micro-CT images show the hypoplasia of the tooth crown consisting of enamel and dentin. Tooth root shortening was also observed in *Slc26a2* deficient tooth germ. Scale bars, 450 μm (C) Quantitative assessment of the dentin volume demonstrated a significant reduction in *Slc26a2-KO-△ exon2* mice tooth germ compared to the control tooth germ. Means ± s.d. (n = 5) are shown as horizontal bars. *P* values were determined by unpaired Student’s *t*-test. \**P* < 0.05.

### Slc26a2-deficient tooth germs showed significantly reduced dentin formation compared to the control tooth germs in *ex vivo* organ culture under the kidney capsule

Due to the neonatal lethality of *Slc26a2-KO-△exon2* mice, it is not possible to assess the effects of *Slc26a2* deficiency on the dentin and enamel matrix production in *Slc26a2-KO-△exon2* mice. We performed *ex vivo* organ culture of tooth germs by implanting the tooth germs under the kidney capsule of nude mice. After four weeks of organ culture, the implanted tooth germs were collected and analyzed by micro-CT. The sagittal sections of the micro-CT images showed hypoplasia of the tooth crown, consisting of enamel and dentin (Fig. 4B). A quantitative assessment of the dentin volume demonstrated a significant reduction in *Slc26a2-KO-△exon2* mouse tooth germ compared to the control tooth germ (Fig. 4B). In addition, tooth root shortening was observed in *Slc26a2-*deficient tooth germs (Fig. 4B). These results might support our assertion that *Slc26a2* deficiency leads to impairment of odontoblast differentiation.

### Sulfate transporter defect due to Slc26a2 deficiency is partly compensated in lower tooth germ

Since the phenotype of *Slc26a2-KO-△exon2* mouse upper tooth germ is more pronounced than that of lower tooth germ (Fig. 3), we hypothesized that the function of *Slc26a2* as a sulfate transporter would be compensated for in the lower tooth germ. To examine the sulfate uptake in the upper and lower tooth germs in *Slc26a2*-deficient and control mice, we quantified the amount of sulfated glycosaminoglycan (GAG), which reflects the sulfate uptake through its transporter. The total amount of sulfated GAG in the upper molar tooth germs was significantly (P<0.0001) decreased by *Slc26a2* deficiency (Fig. 5A). In contrast, there was no significant difference on the total amount of sulfated GAG in the lower molar tooth germs between the *Slc26a2-deficient* and control tooth germs (Fig. 5A). While there was no significant difference in the total amount of sulfated GAG between the upper and lower molar tooth germs of the control group, the total amount of sulfated GAG in *Slc26a2-deficient* tooth germs was significantly decreased in the upper molars compared to the lower molars.

**Figure 5.**
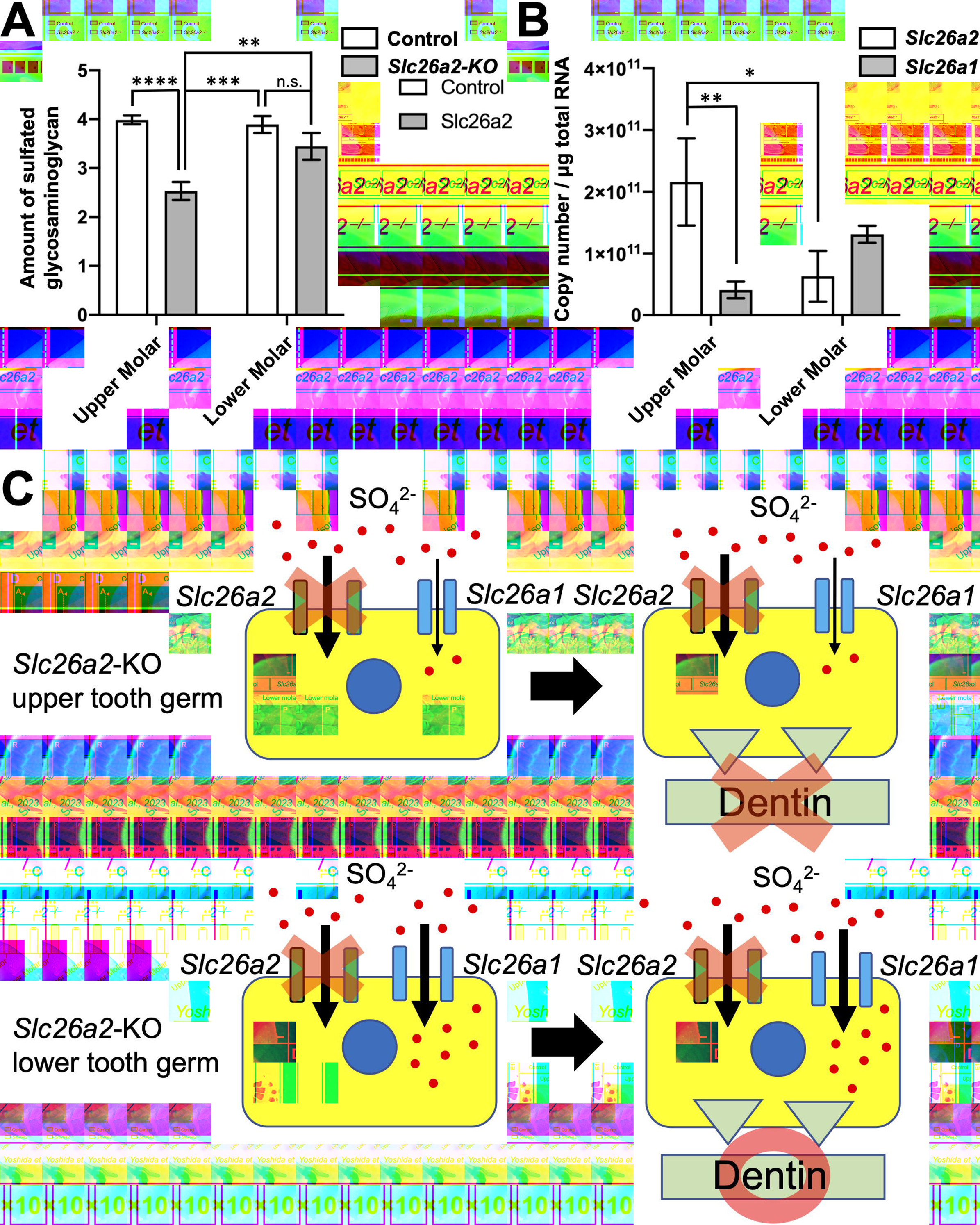
The sulfate transporter defect by *Slc26a2*-deficiency is partly compensated in lower tooth germ. (A) The total amount of sulfated GAGs in upper molar tooth germs was significantly decreased by *Slc26a2* deficiency. Means ± s.d. (n = 3) are shown as horizontal bars. *P* values were determined by two-way ANOVA. \*\**P*<0.01, \*\*\**P*<0.001, \*\*\*\**P*<0.0001. (B) The odontoblasts layer was extracted by micro-dissection from the sagittal section of upper and lower molars. Absolute quantification was performed to evaluate the *Slc26a1* and *Slc26a2* expression. The expression of *Slc26a2* was significantly higher than *Slc26a1* expression in odontoblast of upper tooth germs. Conversely, the expression of *Slc26a1* is higher than *Slc26a2* in odontoblasts of lower tooth germs. Means ± s.d. (n = 3) are shown as horizontal bars. *P* values were determined by two-way ANOVA. \**P*<0.05, \*\**P*<0.01. (C) Schematic illustration of the genetic redundancy between *Slc26a1* and *Slc26a2* in upper and lower tooth germs.

To assess the genetic redundancy by *Slc26a1,* a homolog of *Slc26a2*, and *Slc26a2*, we performed micro-dissection, which enables the extraction of RNA from odontoblasts with high purity and the absolute quantification of mRNA. As a result, we found that the expression of *Slc26a2* was significantly higher than that of *Slc26a1* in odontoblasts from the upper tooth germ (Fig. 5B). Conversely, the expression of *Slc26a1* was higher than that of *Slc26a2* in odontoblasts from the lower tooth germs (Fig. 5B). These results suggest that *Slc26a1* is predominantly expressed in lower tooth germs and may compensate for the function of *Slc26a2* in *Slc26a2*-deficient lower tooth germs (Fig. 5C).

## Discussion

In the present study, we demonstrated that *Slc26a2* is predominantly expressed in dental tissues, including odontoblasts and ameloblasts, during tooth development (Fig. 1). A deficiency of *Slc26a2* in chondrocytes reportedly disrupts cartilage growth via the attenuation of chondrocyte proliferation and induction of cell death (Zheng et al., 2019). We also confirmed that cell proliferation was decreased and apoptosis was increased in *Slc26a2-*deficient chondrocytes *in vivo* (Appendix Figure 4). However, the cell proliferation and apoptosis were not markedly altered in the tooth germ of *Slc26a2-KO-△exon2* mice compared to the control mice (Appendix Figure 5). The sulfation level is markedly higher in cartilage than in the developing tooth germ (Appendix Figure 6). This indicates that the consumption of the sulfate anion in developing tooth germ is much lower than in cartilage, and susceptibility to *Slc26a2* deficiency may be dependent on the tissue-/cell-specific requirement of sulfate transportation. Interestingly, the tooth phenotype in *Slc26a2-KO-△exon2* mice is more prominent in the upper incisors and molars than in the lower teeth (Fig. 3). We found that the expression of *Slc26a1* was higher in the lower tooth than in the upper tooth (Fig. 5B), suggesting that SLC26A1 may be able to compensate for the disability of SLC26A2-mediated sulfate transportation in lower tooth germ (Fig. 5C). In fact, *Slc26a2* deficiency does not affect the amount of sulfated GAG in the lower tooth germ instead of showing a significant difference in the upper tooth germ of *Slc26a2-KO-△exon2* mice compared to the control mice (Fig. 5A). The upper and lower jaws are derived from the first branchial arches (BA). Dlx transcriptional factors are regionally expressed within BAs and are implicated in regulating jaw-specific genetic programs for proper patterning during craniofacial development (Dollé et al., 1992) (Bulfone et al., 1993) (Robinson and Mahon, 1994) (Depew et al., 2002) (Qiu et al., 1997). The *Slc26a1* and *Slc26a2* expression patterns might be part of the jaw-specific gene regulation machinery. Further studies will be required to clarify the mechanisms underlying the regulation of the different expression patterns of *Slc26a1* and *Slc26a2* in the upper tooth germ and lower tooth germ during tooth development.

*Slc26a2-KO-△exon2* mice show short, flattened odontoblasts, implying the loss of the odontoblasts’ polarity (Fig. 3A–D). The expression of *Dspp* and *Dmp1*, well-defined odontoblast differentiation markers, was also decreased in *Slc26a2*-deficient tooth germ *in vivo* (Fig. 4A). These results indicate a critical role for *Slc26a2* in odontoblast differentiation and dentin formation. In addition, the odontoblastic differentiation of dental pulp stem cells was substantially suppressed in *Slc26a2* knockdown cells *in vitro* (Fig. 4C). This result suggests that *Slc26a2* directly affects the odontoblast differentiation instead of applying secondary effects via the surrounding tissue or systemic sulfate insufficiency. Due to the postnatal lethality of *Slc26a2*-deficient mice, it is impossible to evaluate the effects of *Slc26a2* deficiency on the tooth morphogenesis and dentin formation *in vivo*. In the present study, we used *ex vivo* transplantation of tooth germ under the kidney capsule to demonstrate the reduced dentin formation in *Slc26a2*-deficient tooth germ compared to the control tooth germ (Fig. 4D, E). *Ex vivo* organ culture of tooth germ is often selected as an alternative method for examining tooth germs from genetically modified mice. Kidney-capsule grafting has been previously reported to provide tooth germs with an *in vivo* biological environment (Ono et al., 2017; Otsu et al., 2012). It is thus possible to retain the physiological features, and morphogenesis of transplanted tooth germs proceeds normally under conditions of kidney-capsule grafting.

GAGs are linear polysaccharides consisting of repeated disaccharide units and exist as proteoglycans (PGs) by attaching to specific serine residues in the core protein. Sulfation of GAGs produces a negative charge at various positions on PGs and induces diverse biological functions of PGs due to structural microheterogeneity. Heparan sulfate is a highly sulfated GAG involved in odontogenesis and amelogenesis (Bishop et al., 2007). Yasuda et al. generated mice deficient in Golgi-associated N-sulfotransferase 1 (Ndst1), which mediates the sulfation of HS-PG glycosaminoglycan chains. Ndst1 knockout mice show hypodontia of incisor and molar formation and showed the abnormal differentiation and organization of odontoblasts (Yasuda et al., 2010). Hayano et al. reported that removal of the sulfate groups from the 6-*O*-position of *N*-acetylglucosamine by the extracellular glucosamine-6-sulfatases *Sulf1* and *Sulf2* was an important process in the differentiation of odontoblasts and production of the dentin matrix during dentinogenesis. More specifically, *Sulf1/Sulf2* double-null mice showed a thin dentin matrix and short roots, with a reduced *Dspp* mRNA expression (Hayano et al., 2012). Chondroitin sulfate is also indicated to be involved in tooth development (Ida-Yonemochi et al., 2022). Our results regarding the diminished odontoblast differentiation and dentin matrix production in *Slc26a2-KO-△exon2* mice are consistent with the phenotype of transgenic mice with a genetic modification of sulfated GAGs. *Slc26a2* deficiency may thus affect odontogenesis via the reduction of GAG sulfation during tooth development.

In conclusion, our results demonstrate that *Slc26a2* is predominantly expressed in tooth germ among the sulfate transporter gene, and *Slc26a2* deficiency in mice leads to hypoplasia of the incisors and molars, particularly in the upper tooth germ. Odontoblast differentiation and dentin matrix formation are impaired by *Slc26a2* deficiency in tooth germ. This is the first paper to demonstrate the pivotal roles of the SLC26A2-mediated sulfate transporter during tooth development. Our results may provide insight into the mechanisms underlying tooth abnormalities in recessively inherited chondrodysplasia patients caused by mutations in the SLC26A2 gene.

## Author Contributions

1. Y. Yoshida contributed to data acquisition, analysis, and interpretation, and drafted the manuscript; T. Inubushi contributed to conception, design, data acquisition, analysis, and interpretation, and drafted and critically revised the manuscript; M. Yokoyama and P. Nag contributed to acquisition, analysis and interpretation. A. Oka contributed to interpretation of the data and critically revised the manuscript. T. Murotani, R. Kani, Y. Shiraishi contributed to acquisition and analysis; H. Kurosaka contributed to interpretation of the data; Y. Takahata and R. Nishimura contributed to Methodology and provided critical resources; P. Papagerakis contributed to critically revised the manuscript; T. Yamashiro contributed to critically revised the manuscript and conception of the study. All authors gave final approval and agreed to be accountable for all aspects of the work.

## Acknowledgments

We thank Ms. Yuki Okamoto and Mayumi Yoshimoto for the excellent care and maintenance of our mouse colony and for valuable assistance in the histological, molecular and protein work. This work was funded by Grants-in-Aid for Scientific Research (19KK0232 and 20K21677 to T. Inubushi, 22K10242 to A. Oka, 22K21037 to Y. Yoshida) from the Japan Society for the Promotion of Science. The authors declare no potential conflicts of interest with respect to authorship and/or publication of this article.

## Data Sharing Statement

The data that support the findings of this study are available from the corresponding author upon reasonable request.

