## Appendix file for "SLC26A2-mediated sulfate metabolism is essential for the tooth development"

### Appendix Materials and methods

**Ethics statement**

All animal experiments were performed in strict accordance with the guidelines of the Animal Care and Use Committee of the Osaka University Graduate School of Dentistry, Osaka, Japan. The protocol was approved by the Committee on the Ethics of Animal Experiments of Osaka University Graduate School of Dentistry. Welfare guidelines and procedures were performed with the approval of the Osaka University Graduate School of Dentistry Animal Committee. Approval number: 3745, 29-033-0.

**Animal**

Embryos were washed twice with Opti-MEM solution and aligned in the electrode gap filled with 50 µl of Cas9/gRNA (crRNA-tracrRNA complex)/ssODN (200/100/100 ng/µl) mixture. The intact embryos were subjected to electroporation using poring (225V) and transfer pulses (20V). After electroporation, embryos were returned to KSOM Mouse Embryo Media (Millipore Sigma) at 37°C. Genome-edited 2-cell embryos were transferred to pseudopregnant ICR female mice oviducts, and genomic DNA from newborn mice was analyzed by PCR. The sequence of gRNA is left gRNA: 5'-AGTCTGAGACCGGTCATGGC-3' and right gRNA: 5'-ACAATGA GCTCGACCGGAAT-3'. In all experiments, *Slc26a2*$^{wild/\mathrm{wild}}$ offspring were used as controls. Genotyping of mice and embryos was performed by PCR with the specific primers listed in supplemental table 1. Mice were fed ad libitum on solid feed and sterile water irradiated with ultraviolet light. The environmental conditions of the animal facility were maintained at constant temperature and humidity and kept under a 12-hour light-dark cycle (8:00-20:00 as the light period).

**Tissue preparation, histology and *in situ* hybridization**

The digoxigenin-labeled RNA probes used in this study were prepared using a DIG RNA labeling kit (Roche) according to the manufacturer's protocol using each cDNA clone as the template. The probes were synthesized from fragments of *Slc26a2* (Allen Institute for Brain Science) and were amplified with T7 and SP6 adaptor primers through a PCR. After hybridization, the expression patterns for each mRNA were detected and visualized according to their immunoreactivity with anti-digoxigenin alkaline phosphatase-conjugated Fab fragments (Roche). A minimum of three embryos of each specimen type were examined per probe.

**Immunohistochemistry and TUNEL staining**

Immunofluorescence staining was accomplished overnight at 4°C on 15 μm sections using polyclonal rabbit anti-Ki67 (1:400, ab15580, Abcam). Sections were then counterstained with DAPI (1:500, Dojindo) and mounted using fluorescent mounting media (Dako). At least three embryos were used for each genotype for each analysis. According to the manufacturer's instructions, apoptotic cells were identified using an in situ cell death detection kit (11684795910, Roche).

**Whole-mount skeletal staining**

Mice were fixed in 95% ethanol overnight at room temperature. They were then left in acetone overnight at room temperature and incubated overnight in a cartilage staining solution containing 0.03% (w/v) Alcian blue, 80% ethanol, and 20% acetic acid. The first rinse was performed with several changes of 70% ethanol. To improve visibility of cartilage morphology, washings were terminated before cartilage was completely de-stained. Ossified tissue was stained with an alizarin red solution containing 0.005% (w/v) alizarin red in 1% (w/v) KOH for 4 hours at room temperature and sequentially placed overnight at 4°C to delay staining. Samples were placed in a 50% glycerol solution containing 1% (w/v) KOH to remove excess staining.

**Micro-Computed Tomography (MicroCT)**

Maxilla and mandible were collected from control and *Slc26a2-KO-△exon2* mice at P0. These tissues were fixed in 4% paraformaldehyde overnight. Embryos are placed in 70% ethanol for 1 day and 100% ethanol for 3 days. after that embryos are Placed in 100% ethanol with xylene(1:3) for 1 hour. Then place embryos in 100% ethanol, 90%, 80%, and 70% for 30 minutes each; place in 70% ethanol containing 1% phosphotungstic acid for 1 week for contrast.Both maxilla and mandible were then scanned by micro-CT (R_mCT2, Rigaku) at 90KV, 200μA, microfocus 5μm/voxel size. Volume Graphics (VGstudio) MAX 2.2 software was using for reconstruction of three-dimensional images.

**Laser microdissection**

Dissected heads were freshly mounted in Tissue-Tek and immediately frozen. The tissue was then sectioned serially at a thickness of 20 μm using a cryostat (Leica CM 1950). Sections were mounted on film-coated slides; whole sections were obtained consecutively from the anterior palate at E18.5 and stained with Hematoxylin and Eosin (HE). Odontoblasts from maxillary and mandibular molar were collected in tubes and separated from the sample sections with a manual laser capture microdissection system (LMD6500, Leica). Tissues were serially sectioned at -20°C with a thickness of 25 μm using a cryostat (CM 1950, Leica).

**RNA extraction and qRT-PCR analysis**

The protocol for RNA extraction and qPCR analysis was as reported previously (Inubushi et al. 2012). Total RNA was extracted from dissected tissue following the manufacturer's protocol using RNeasy mini kit (Qiagen), with purity and quantity assessed by Nanodrop spectrophotometer. The extracted RNA was reverse transcribed to cDNA using an oligo dt with reverse transcriptase (Takara bio). For real-time PCR, aliquots of total cDNA were amplified using Fast SYBR Green PCR Master Mix (Applied Biosystems, Foster City, CA, USA) or Fast TaqMan Fast Universal PCR Master Mix (Applied Biosystems, Foster City, CA, USA). Data were acquired and analyzed with the Step One Real-Time PCR System using Step One Software, Version 2.1 (Applied Biosystems). The PCR products were quantified with Gapdh as the reference gene. The primers and probes were listed in Appendix 1 Table 2. Each experiment was performed in triplicate.

**Reanalysis of public scRNA-seq data**

The public scRNA-seq data set for GSE146855 of mice incisors was downloaded from the GEO database to reanalyze the expression profile in odontoblast differentiation. The data was analyzed using a package of Seurat (version: 4.0.5) with R studio. A total of 6,260 cells were reanalyzed. After normalization, scaling, and principal component analysis (PCA) of the data, Cells were clustered using FindNeighbors (dims = 1:6) followed by FindClusters (resolution = 0.35). The RunUMAP function was used to visualize the cell clusters. Differential expression and cell identification were performed using FindAllMarkers (min.pct = 0.25, logfc.threshold = 0.25) with the Wilcoxon rank sum test. Top 5 differentially expressed features (cluster biomarkers) and the cluster biomarkers are shown in appendix figure 1. Visualization of *Slc26a1, Slc26a2*, *Slc26a6*, *Slc26a7*, *Slc26a10*, and *Slc26a11* gene expressions with dot plot was generated with Seurat function DotPlot.

**Cell culture and lentivirus transduction**

Human Dental pulp stem cells (hDPSCs) isolated from human adult third molars (Lonza) were cultured in Dulbecco's Modified Eagle's Medium (Wako) containing 20% fetal bovine serum (Invitrogen) and 1% penicillin/streptomycin (Sigma-Aldrich), which was designated as growth medium (GM). To induce odontoblasts differentiation, hDPSCs were cultured in odontoblasts differentiation medium (EM) consisting of alpha Modified Eagle's Medium (Wako) containing 20% fetal bovine serum (Invitrogen) supplemented with 10 nM dexamethasone, 10 mM β-glycerophosphate, and 50 μg/mL vitamin C (Sigma-Aldrich). To knockdown *Slc26a2* expression in hDPSCs, we used lentivirus-mediated shRNA transduction. Lentivirus particles expressing an shRNA that is validated to deplete human *Slc26a2* (Mission shRNA, TRC Clone ID: TRCN8607, MilliporeSigma) and control lentivirus particles expressing an shRNA that does not target any known genes (Mission shRNA, MilliporeSigma SHC005) were purchased from MilliporeSigma. Lentivirus particles were added to hDPSCs cultured in growth media supplemented with 5 μg/ml polybrene and cultured for 2 days. Cells transduced with lentiviral shRNAs were selected and maintained in the presence of 10 μg/mL puromycin.

**Blyscan sulfated glycosaminoglycan assay.**

Briefly, $4 \times 10^5$ cells were plated in a T25 flask, and 48 h later, the cells were treated with 1 μg/ml DS-500 or 50 mM NaClO3. Twenty-four hours after treatment, the medium was aspirated, and the cells were rinsed with phosphate-buffered saline (PBS). The cells were lysed in papain extraction reagent added to the cell monolayer for 3 h at 65°C. The total cell extract containing total GAGs was harvested, and samples were centrifuged at 10,000 × g for 10 min. A total of 100 µl of the supernatant was used for the assay.

**Alcian blue staining**

Stain with Alcian blue for 30 minutes, and rinse with tap water for 5 minutes. Rinse with tap water for 1 minute, dehydrated with ethanol, washed with xylene, and mount with admixture.

**Statistical analysis**

Statistical methods were not used to predetermine sample size. Statistical analyses were performed with GraphPad Prism 8. Students' two-tailed t-tests and two-way ANOVA were used under the assumption of normal distribution and observance of similar variance. $P<0.05$ was considered significant. Bonferroni post hoc analysis was performed where applicable. Values are expressed as mean±s.d. For all of these experiments, variances between groups were similar and data were symmetrically distributed. Data shown are representative images; each analysis was performed on at least three mice per genotype. Immunostaining was performed at least in triplicate. For other experiments, the numbers of biological replicates, animals, or cells are indicated in the text. No randomization was used to allocate experimental units. There is no exclusion and inclusion criteria, all mutant mice were allocated to each experiment. There were no excluding animals. Confounders were not controlled in the experiments. For the quantitative measurements and statistical analysis, genotype information was blinded.

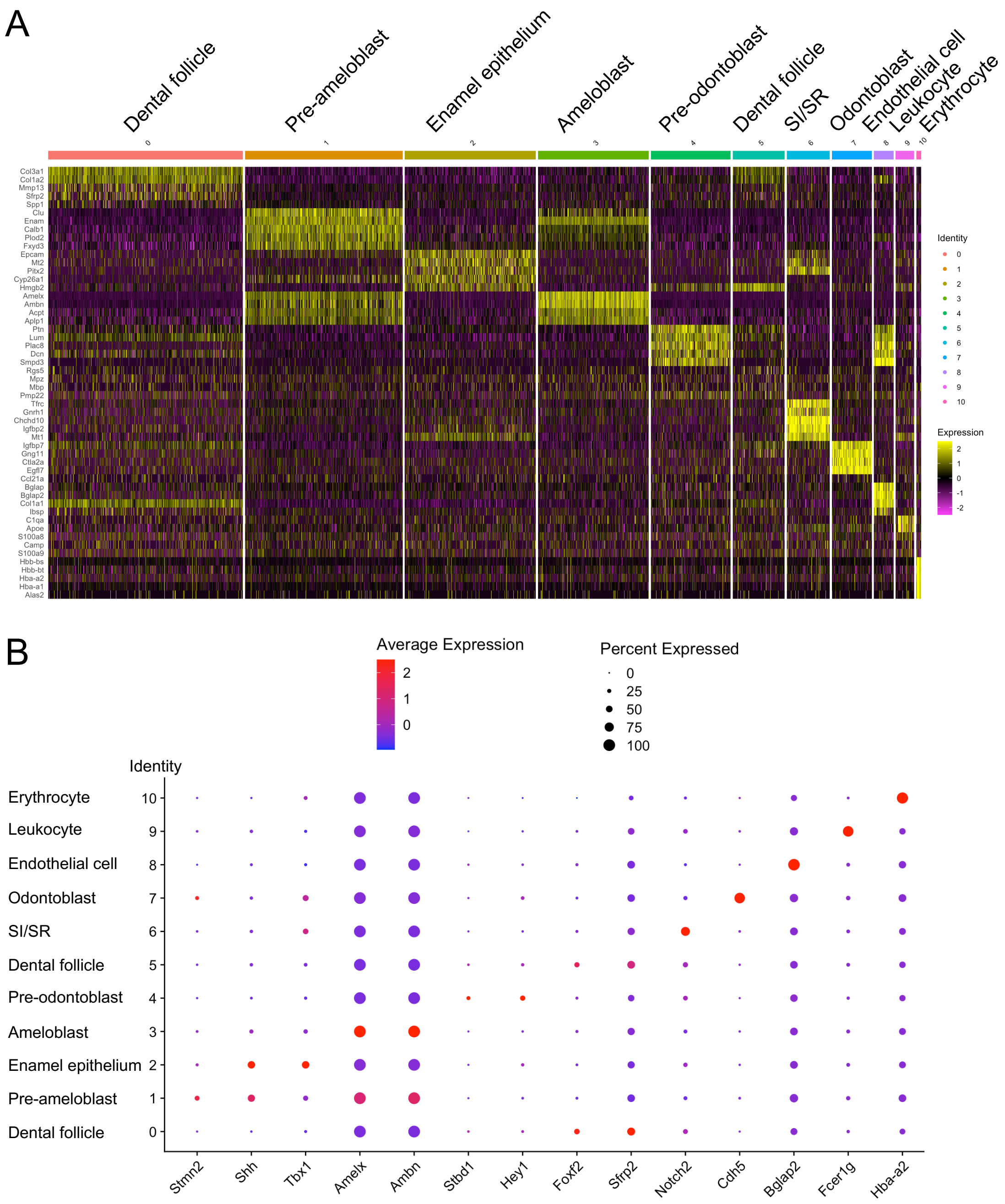


**Appendix Figure 1. Reanalysis of the public scRNA-seq dataset (GSE146855) of the isolated mice incisors.**

(A) Visualization of top 5 differentially expressed features (cluster biomarkers) with Seurat function DimHeatplot. (B) The cluster biomarkers are shown with Seurat function Dotplot.

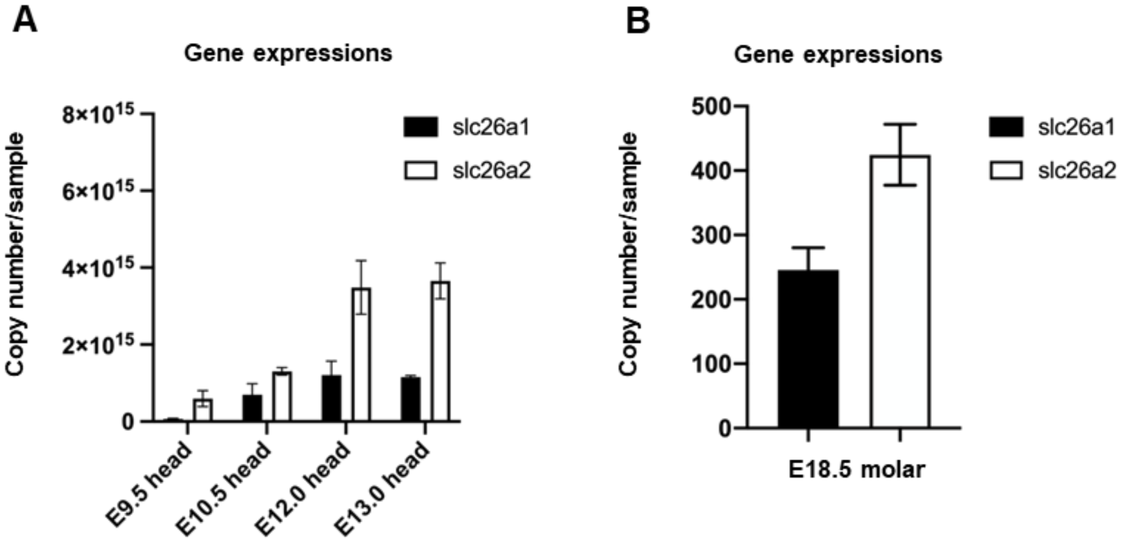


**Appendix Figure 2. Expression patterns of *Slc26a1 and Slc26a2* in craniofacial development.** (A) The Expression pattern of *Slc26a1 and Slc26a2* gene during normal development of the craniofacial region. Total RNAs were extracted from the craniofacial region at E9.5-E13.0. At each developmental stage, the expression of *Slc26a2* was found to be markedly higher than that of *Slc26a1*. Furthermore, the expression of *Slc26a2* increased with developmental stage, with craniofacial formation beginning at E9.5 and the tooth bud forming at E13.5 (B) The expression pattern of *Slc26a1* and *Slc26a2* during normal upper and lower molars development at E18.5. The expression of *Slc26a1* and *Slc26a2* was evaluated by qPCR. The expression level of *Slc26a2* of the upper molars was found to be higher than that of the lower molars. Gapdh was used as an internal control for normalization. Means ± s.d. (n=3) are shown as horizontal bars. **P<0.01 (two-way ANOVA).

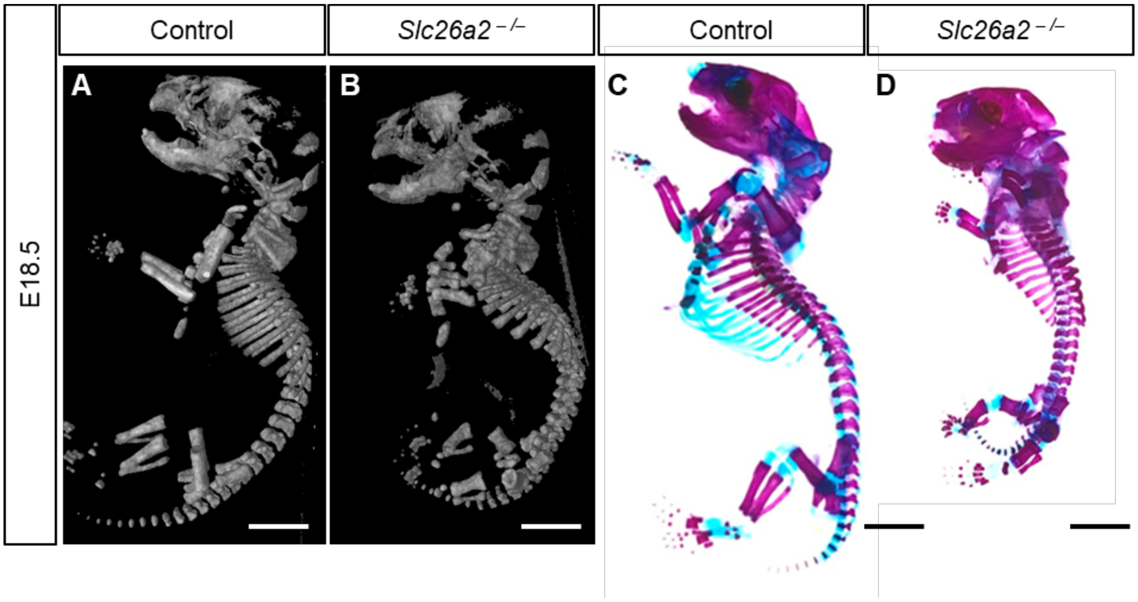


**Appendix Figure 3. Morphological analysis of *Slc26a2-KO-Δexon2* at E18.5.**

(A) Micro-CT images demonstrate a short stature, small chest, and very short limbs in *Slc26a2-KO-Δexon2* compared to the control mice. The long tubular bone is shorter in length and longer in diameter than the control mice. (B) Whole-mount skeletal preparations showed that *Slc26a2-KO-Δexon2* had chondrodysplasia and reduced alcian blue staining of the cartilage compared to the control mice at E18.5. Scale bars, 4mm. Data shown are representative images; each analysis was performed on at least three mice per genotype.

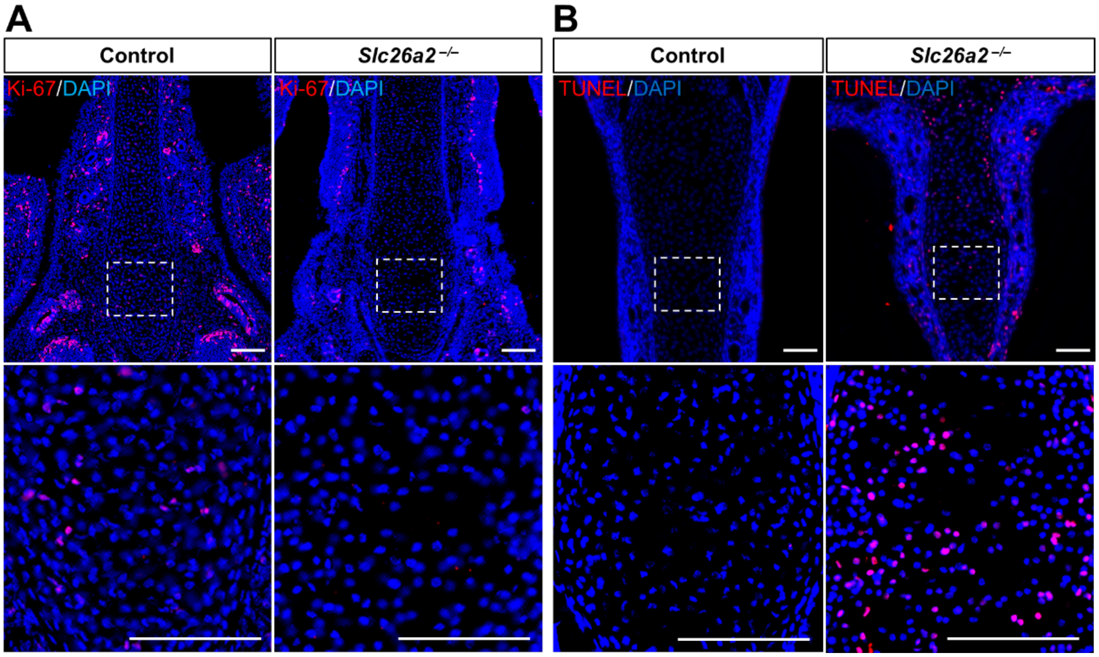


**Appendix Figure 4. Deficiency of *Slc26a2* in nasal septal cartilage chondrocytes decreased cell proliferation and increased apoptosis.**

(A) Frontal sections of maxillary from control and *Slc26a2-KO-Δexon2* embryos were immuno-labelled with anti-Ki67 antibody. Analysis of cell proliferation in nasal septal cartilage chondrocytes of E18.5 embryos. The number of Ki67 positive cells were decreased in *Slc26a2-KO-Δexon2* compared to the control embryos. The lower panels are the enlarged images of the boxed area in the upper panels. (B) Analysis of apoptosis in nasal septal cartilage chondrocytes of E18.5 embryos. Apoptotic cells were detected by TUNEL assay. The number of TUNEL positive cells were decreased in *Slc26a2-KO-Δexon2* compared to the control embryos. The lower panels are the enlarged images of the boxed area in the upper panels. Scale bars, 50 µm in A-B. Data shown are representative images; each analysis was performed on at least three mice per genotype.

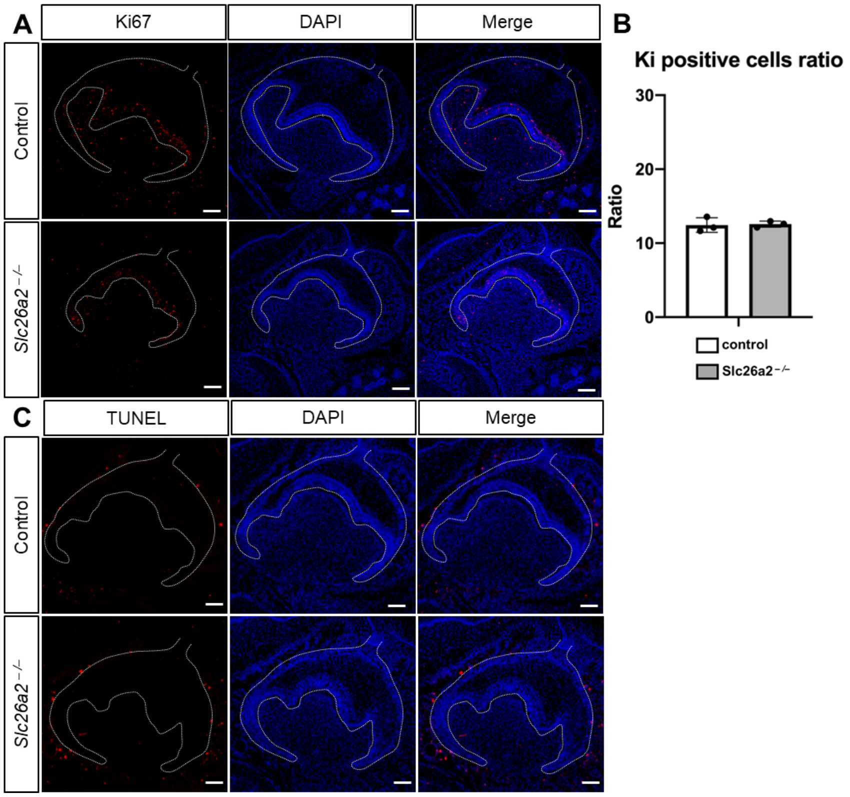


**Appendix Figure 5. The cell proliferation and apoptosis were not altered in the tooth germ of *Slc26a2-KO-△exon2* compared to the control mice.**

(A) Analysis of cell proliferation in the tooth germ in *Slc26a2-KO-△exon2* and control embryos at E18.5. Frontal sections of the upper tooth germ were immuno-labeled with anti-Ki67 antibody. (B) Quantitative analysis of the number of Ki67 positive cells in (A). Means ± s.d. (n=3) are shown as horizontal bars. *P* values were determined by unpaired Student's *t*-test. There is no statistical difference in the number of Ki67 positive cells between *Slc26a2-KO-△exon2* and control embryos. (C) Analysis of apoptosis in the tooth germ of E18.5 embryos. Frontal sections of the tooth germ of E18.5 embryos were analyzed by TUNEL assays. TUNEL positive cells in dental epithelium and mesenchyme were not detected in both *Slc26a2-KO-△exon2* and control embryos. Scale bars, 50 μm. Data shown are representative images; each analysis was performed on at least three mice per genotype.

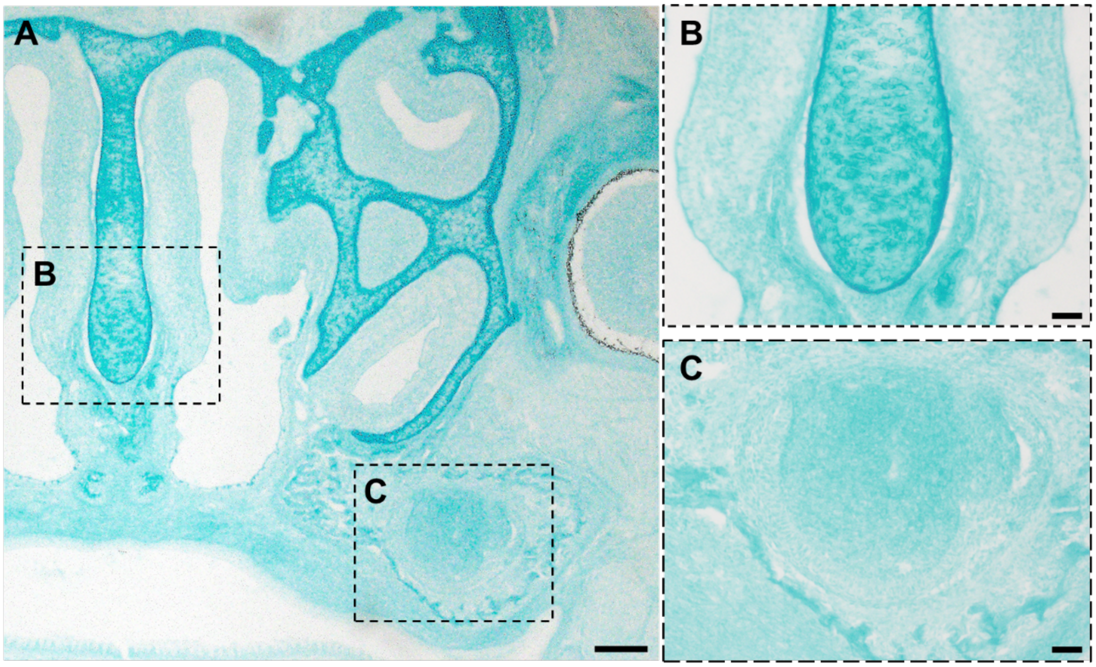


**Appendix Figure 6. The sulfation level is extremely higher in cartilage compared to the developing tooth germ.**

(A) Frontal sections of the maxillary including the nasal cartilage (B) and the upper molar (C) in wild type mice were stained with alcian blue dye at pH2.5. Acidic polysaccharides mainly composed of sulfated glycosaminoglycan were colored with blue. Right panels are enlarged images of the boxed area. The intensity of the alcian blue is higher in the nasal cartilage compared to the developing tooth germ. Scale bars, 100µm in A; 50µm in B, C. Data shown are representative images; each analysis was performed on at least three mice per genotype.

Appendix Table1. Primer sequences for PCR genotyping of mice

| | |
|---|---|
| *Slc26a2 mutant* forward | 5’-AAGCCTTTGGTTTCCCATCTGA-3’ |
| *Slc26a2 mutant* reverse | 5’-TGGGAATGTGTCCAGCTTAATCG-3’ |
| *Slc26a2 wild* forward | 5’-TTGAGGGCCATCATTTTAGCAGC-3’ |
| *Slc26a2 wild* reverse | 5’-CCAGCTATTCTTCCCCTTCCTCTC-3’ |

Appendix Table2. Primers used for qRT-PCR

| | |
|---|---|
| SLC26A2- forward | 5′-CCA GAT GTG GAG GAT TAG CAG AAT GG-3′ |
| SLC26A2- reverse | 5′-ACA GCT TCA TAA TCT CTG CGA ACT TCT TTC AGT GT-3′ |
| DSPP-forward | 5’-TGCATTTGGGCAGTAGCAT-3’ |
| DSPP- reverse | 5’-TGTCTCTCCAGTGGTTTGCTT-3’ |
| DMP1-forward | 5’-CCCTTGGAGAGCAGTGAGTC-3’ |
| DMP1- reverse | 5’-CTCCTTTTCCTGTGCTCCTG-3’ |
